# Epithelial function of the circadian clock gene, *Bmal1*, in regulating the mucosa

**DOI:** 10.64898/2026.04.15.718752

**Authors:** Z. Taleb, C. Edwards, R. Wan, M. Fatmah, M. Haireek, H. Wang, W.I. Khan, P. Karpowicz

**Affiliations:** Department of Biomedical Sciences, University of Windsor, Windsor, ON, Canada; Department of Pathology and Molecular Medicine, McMaster University, Hamilton, ON, Canada; Farncombe Family Digestive Health Research Institute, McMaster University, Hamilton, ON, Canada

**Keywords:** Circadian rhythms, colitis, mucus, regeneration

## Abstract

Circadian rhythms, 24-hour repeating oscillations in daily physiology, are implicated in maintaining intestinal homeostasis. These rhythms are driven by the circadian clock, a molecular timekeeper found throughout cells of the body, including those of the intestinal epithelium. Loss of clock function has been found to worsen colitis; however, it is not clear how the clock impacts regeneration which enables a tissue to return to its homeostatic set point following an injury. To investigate these questions, we used a conditional knockout of the core clock gene, *Bmal1*, in mouse colon epithelial cells. Our data show that prior to injury *Bmal1* promotes colon mucus production, which increases in thickness and within goblet cells when mice are active and begin feeding. *Bmal1* loss lowers mucus production but does not drive an apparent tissue phenotype until the system is injured and regenerates itself. In this context, *Bmal1* epithelial loss drives a male-specific colitis phenotype and a delay in the ability of colon epithelial cells of both male and female mice to resolve injury to return to their homeostatic set point. Our data suggest that epithelial sex-specific clock rhythms are needed for optimal colon barrier homeostasis.

## Introduction

The circadian clock is a molecular system that drives 24-hour rhythms in physiology (1). Clocks located in the cells of the body are thought to organize and coordinate daily cellular timing with the 24-hour daily environment. This includes cells of the gastrointestinal tract where circadian rhythms in digestion, motility, immune function, and cellular proliferation have been documented (2–5). The core clock is a transcriptional timer that comprises the E-Box binding transcription factors Clock and Bmal1 which heterodimerize to drive the expression of thousands of genes including their negative regulators, *Per* and *Cry* (1). This transcriptional feedback system provides 24-hour rhythms in gene expression; regulators including *Nr1d1/2* and the *Rorα* family provide additional feedback systems to maintain 24-hour rhythmicity. Clock-driven transcription rhythms promote optimal tissue function: the disruption of the circadian system, through shift-work and other factors, has been implicated in many pathologies including Inflammatory Bowel Disease (IBD) (5–9). The clock system, which includes rhythms in behaviour, cardiovascular function, body temperature, and metabolism is distributed throughout the body and coordinated through brain-driven and hormone-driven processes (10). In short, a combination of system-wide circadian rhythms with rhythms driven by intestinal cell clocks are a fundamental feature of gastrointestinal physiology.

Gastrointestinal tissues are protected by an epithelial barrier that separates the gut lumen and the body. The epithelium is organized into crypts where intestinal stem cells continuously replenish epithelial cells including enterocytes / colonocytes, goblet cells, and enteroendocrine cells (11). The epithelium is also known as the mucosa, because its numerous goblet cells produce a protective mucus layer that prevents the gut’s abundant microbiota from contacting gastrointestinal tissue (12). Mouse-based studies have shown that colitis, a form of IBD, arises when the mucus layer separating the bacteria from the cells of the body is reduced (13, 14). Colitis itself is a disease of injury and recovery, where inflammation produces areas of damage in the epithelium, killing epithelial cells that must be regenerated to restore the tissue to its previous state (15–18). The epithelial mucosa is thus a homeostatic system able to return to its original state following inflammatory injury.

Circadian clock function has been previously implicated in colitis. Patients with IBD have been shown to have disrupted expression of circadian clock genes (7), and physiological loss of circadian rhythms in cardiovascular output has been shown to precede impending IBD flares (19). Disrupted circadian rhythms are themselves associated with worse disease outcomes in IBD patients (9, 20). These are mirrored and confirmed in laboratory experiments. Mouse-based studies testing circadian clock genes, including mutants of *Bmal1*, *Per1/2*, *Nr1d1*, and *Rorα* have shown that the loss of these genes leads to an increased severity of colitis (21–25). Of note, *Bmal1* null mutant and *Per1/2* double mutant mice, that lose their 24-hour rhythmicity, also exhibit reduced epithelial cell proliferation and regeneration that is necessary to recover from colitis (21, 23).

*Bmal1* is a non-redundant circadian clock component: its loss completely disrupts clock rhythms (26). A conditional *Bmal1* loss of function allele (27) has been used as a tool to examine the specific contribution of the epithelial circadian clock in colitis. Dextran Sulfate Sodium (DSS) is a chemical model of colitis that mimics the inflammatory injuries present in IBD (28). Using a *TS4-cre* (*Fatty acid binding protein 1* driver), a 2023 study initially showed that the loss of *Bmal1* in the colon epithelium increased DSS-colitis (29). Disease was accompanied by the disruption of epithelial cell adhesion, an inflammatory response, and changes to microbiome composition. In 2024, using a *Villin-cre* epithelial driver, a separate group subsequently found that conditional *Bmal1* loss also increases DSS-colitis (30), and the authors proposed that the epithelial clock regulates a response to the microbiome necessary for circadian clock rhythmicity (31). However, a subsequent 2025 study did not find colitis to be increased in the same *Villin-cre* driven *Bmal1* loss of function even though inflammation / microbiome responses were disrupted (32). Finally, another 2025 study found epithelial *Bmal1* loss to actually decrease DSS-colitis due to *Bmal1* transcription of pro-apoptotic genes whose reduction (in a *Villin-creERT2* + *Bmal1* conditional loss of function) conferred a protective effect through cell survival (33). A discordance in the outcomes of these studies suggests that additional factors may contribute to the effects of epithelial *Bmal1* in homeostasis and/or colitis. Of note, goblet cell number was found to be both lower (30) or unaffected (32) in a conditional *Villin-cre* + *Bmal1* epithelial loss of function.

Discrepancies in the outcomes of *Bmal1* loss in the epithelium may be due to phenotypes that have not been noted in mice prior to the induction of colitis. It is also possible that a sex-specific response is present, except for one of these reports (29), experiments in this area have been performed using only male mice. Finally, while colitis has been tested, it is not clear how the loss of *Bmal1* affects tissue regeneration and recovery to the original state following a colitis injury. To investigate these questions, we re-examined the loss of *Bmal1* in colon epithelial cells. We induced colitis using DSS and assessed epithelial recovery 1 month post-injury. Our findings indicate that loss of *Bmal1* in intestinal epithelial cells impacts mucus production. Goblet cells have transcript rhythms in many genes involved in mucus production and mucus glycosylation, and mucus layer thickness itself exhibits a daily rhythm, driven in part by epithelial *Bmal1* function. We find that male mice exhibit a sex-specific susceptibility to DSS-colitis, but that both male and female mice exhibit a delay in tissue regeneration following injury. These results highlight a basic feature of the mucosal barrier, regulated by the clock, that exists to separate the microbiome and immune system. Disruptions to this process are likely to underlie the effects of *Bmal1* loss on the maintenance of colon tissue homeostasis, thereby explaining some of the discrepancies and downstream mechanisms observed in previous studies.

## Methods

### Animal housing and breeding

*Vil^Cre/+^* (Jackson Laboratories, B6.SJL-Tg(Vil-cre)997Gum/J #004586), *Bmal1^flox/flox^* mice (Jackson Laboratories, B6.129S4(Cg)-Arntltm1Weit/J #007668), and *TdTom* (Ai9, Jackson Labratories, B6.Cg-Gt(ROSA)26Sortm9(CAG-tdTomato)Hze/J #007909) were housed in a barrier facility under a 12-hour light/ 12-hour dark (LD) cycle (with lights on at 7am / Zeitgeber Time 0, and light off at 7pm / Zeitgeber Time 12). Litter mates were used in experiments and were co-housed prior to experimentation. Normal chow was administered *ad libitum*. All mouse procedures were reviewed and approved by animal care regulatory boards at the University of Windsor (#AUPP 23-17).

### DSS colitis / injury

Mice, at 12-14 weeks of age, were housed individually for 1 week before beginning the experiments to acclimate to the new cage. Bedding from their previous co-housed cages was transferred into the new individual cages in order to further control for microbiome effects. Untreated mice were collected following the 1 week acclimation period. To administer DSS, a 2.5% (w/v) solution of DSS (MW: 40 KDa, BOC Sciences, #9011-18-1), dissolved in normal drinking water, was administered *ad libitum* to the mice over the course of 7 days. Consumption of DSS water was measured daily at ZT0 (no differences were observed between controls and mutants in the amount consumed). After a 7-day treatment of 2.5% DSS, the DSS solution was removed and replaced with normal drinking water. Weight and disease activity was monitored daily. If weight loss exceeded 25% over the duration of 3 days, or the mice showed signs of extreme sickness the experiment was terminated and the mice humanely euthanized.

### ClockLab Behavioural Analysis

Mice were housed in an ActiMetrics Circadian Cabinet maintained at 12-hour light/ 12-hour dark (LD) cycle (with lights on at 7am / Zeitgeber Time 0, and light off at 7pm / Zeitgeber Time 12). Locomotor activity was recorded continuously for three weeks, with the first week designated as an acclimation period. Data shown are from the subsequent two weeks. Activity profiles were analyzed using ClockLab Analysis 6, with the final two weeks of recordings normalized and analyzed in 10-minute bins for imaging and 60-minute bins fit to least square change for graphical representation.

### Disease activity

Disease Activity Index (DAI) score included daily measurements of weight loss compared to initial weight, stool consistency, and rectal bleeding. Each disease criteria is assigned a score based on the severity of the symptom (weight loss 0:<1%, 1: 1–5%, 2: 5–10%, 3: 11-14%, 4:>15%, stool consistency 0: normal, 2: loose stools, 4: diarrhea, stool blood 0: negative, 2: positive, rectal bleeding 4: if present) (28).

### Tissue collection

Mice were anesthetized using isoflurane then euthanized using CO_2_ followed by cervical dislocation. For Figures 1-3, colon tissue of untreated animals was collected, measured and immediately placed in ice cold Carnoy’s solution (60% absolute ethanol, 30% chloroform, 10% glacial acetic acid) and fixed for 2 hours. Colons were transferred to absolute ethanol overnight (O/N) before being embedded in paraffin. In the remaining Figures, colon tissue was dissected, measured, and flushed with ice cold phosphate buffer saline (PBS). The tissue was cut longitudinally to open the lumen and was then rolled up from the proximal to distal end (Swiss rolls). The dissected colon tissue was fixed in 4% paraformaldehyde (PFA) (Electron Microscopy Sciences, Hatfield, PA) diluted in PBS for 2 hours.

**Figure 1:**
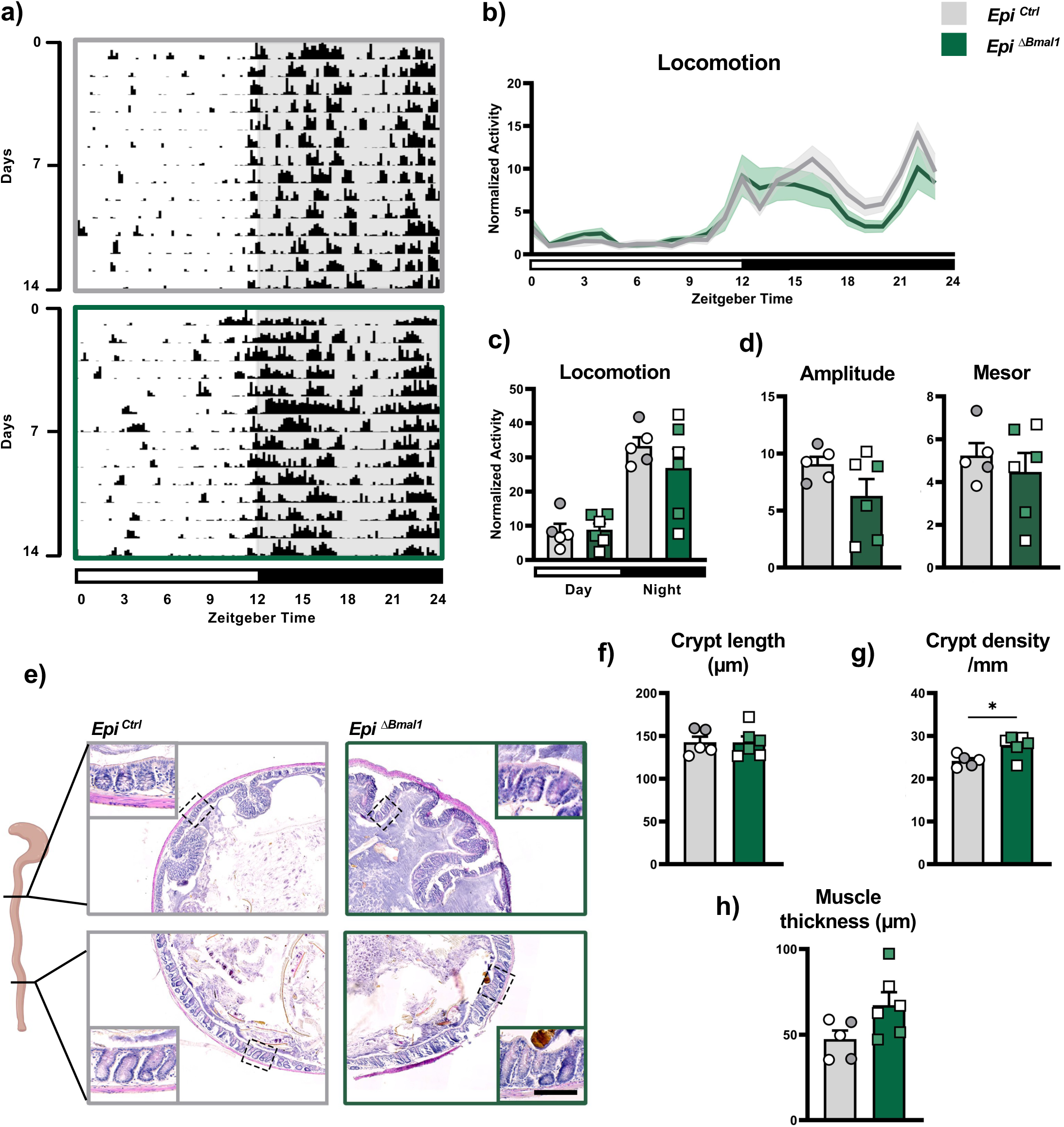
Generation of mice lacking an intestinal epithelial clock. (**a**) Representative actogram of locomotor activity for *Epi^Ctrl^*(grey) and *Epi^ΔBmal1^* mice (green). Lights were maintained at 12:12 LD schedule (indicated in white and light grey) and each row represents a 24-hour day. Individual activity is marked with black traces. (**b**) Average of daily locomotor activity for 14 days in *Epi^Ctrl^* and *Epi^ΔBmal1^* mice (no significant differences). (**c**) Average of daily locomotion, across 12 hours light and 12 hours dark (no significant differences). (**d**) Average amplitude and mesor of daily locomotor rhythms (no significant differences). (**e**) Representative images of Hematoxylin & Eosin (H&E) stained colon transverse sections from *Epi^Ctrl^* and *Epi^ΔBmal1^* mice as indicated in schematic. Magnified inset indicated by a dotted outline, highlighting the selected region for detailed visualization (scale bar= 50μm). (**f**) Distal crypt length (no significant differences). (**g**) Crypt density (# average crypts per mm) shows a significant increase in density in the *Epi^ΔBmal1^* colon. (Unpaired T-test, *p ≤ 0.05). (**h**) Muscle thickness (no significant differences). In the graphs, male mice are indicated in white symbols and female mice as coloured (grey or green) symbols. Full statistical details are provided in Supplementary Table 1.

**Figure 2:**
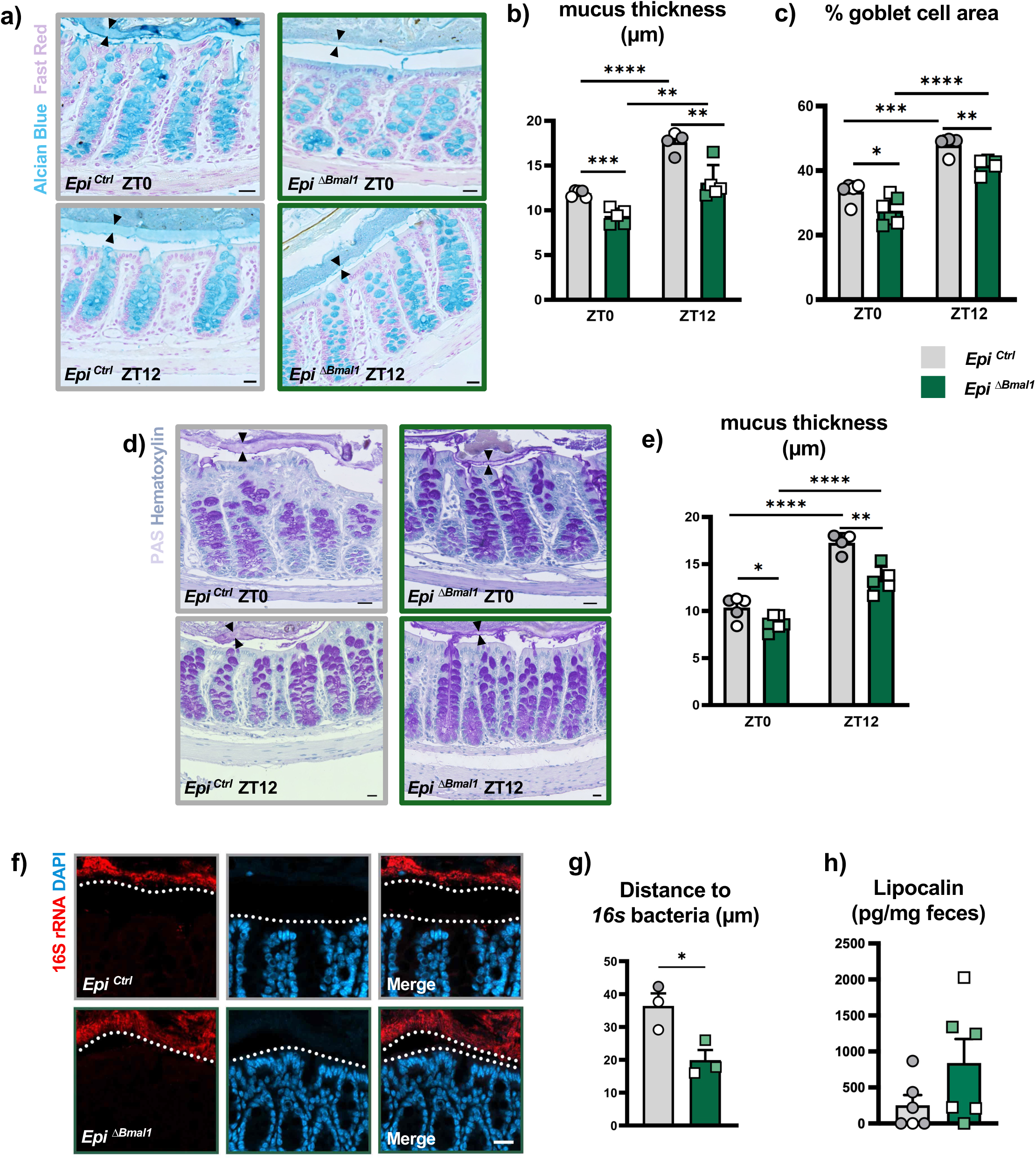
Epithelial loss of *Bmal1* reduces mucus barrier production. (**a**) Representative images of *Epi^Ctrl^* and *Epi^ΔBmal1^* distal colon stained for Alcian blue to label acidic mucus. Shown are samples collected at ZT0 (morning, mice become inactive), and ZT12 (evening, mice become active). Arrowheads indicate mucus thickness. (Scale bar = 20μm). (**b**) Mucus thickness in the distal colon is significantly higher at ZT12 than at ZT0 in *Epi^Ctrl^* but lower at both times in *Epi^ΔBmal1^*(*p ≤ 0.05). (c) goblet cell area in distal colon crypts, defined by the area covered by Alcian blue, shows a similar pattern of change between ZT0 and ZT12 in *Epi^Ctrl^*, and between *Epi^Ctrl^* and *Epi^ΔBmal1^* (*p ≤ 0.05). (**d**) Representative images of distal colon stained for PAS to label neutral + acidic mucus. Arrowheads indicate mucus thickness. (Scale bar = 20μm). (e) Similar to Alcian blue stained mucus, the thickness is higher at ZT12 than ZT0 in *Epi^Ctrl^*, and lower at both times in *Epi^ΔBmal1^*(*p ≤ 0.05). (**f**) Representative images of 16S ribosomal RNA (rRNA) staining of *Epi^Ctrl^*and *Epi^ΔBmal1^* distal colons at ZT0. The dashed line indicates the distance between the 16S+ luminal bacteria and the epithelial cells. (Scale bar = 10μm). (**g**) Quantification of 16S RNA-FISH (at ZT0) indicates significant decrease in bacterial distance in *Epi^ΔBmal1^*(*p ≤ 0.05). (**h**) Lipcalin levels are not significantly different between *Epi^Ctrl^* and *Epi^ΔBmal1^* (p=0.06). In the graphs, male mice are indicated in white symbols and female mice as coloured (grey or green) symbols. Full statistical details are provided in Supplementary Table 1.

**Figure 3:**
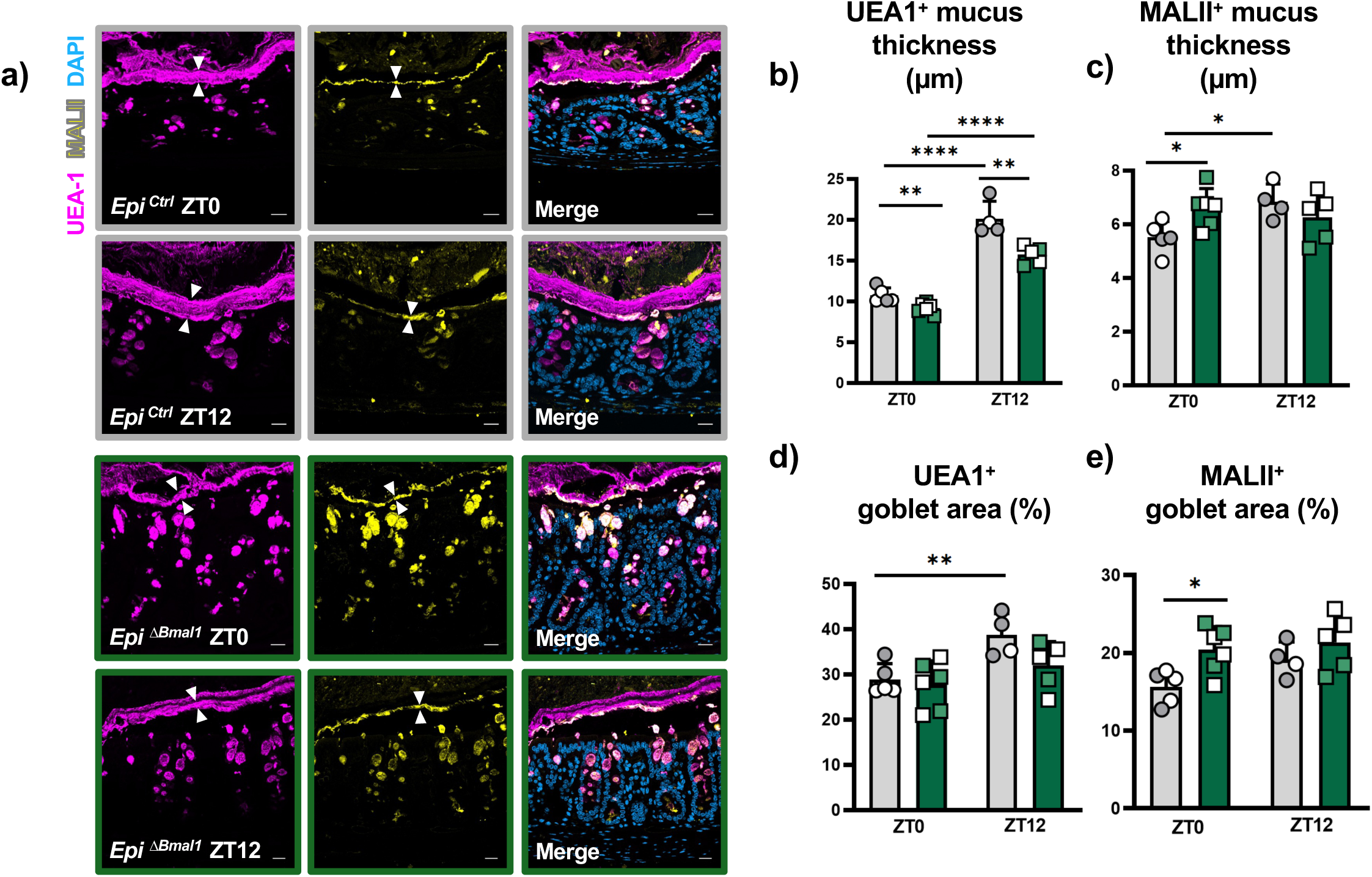
Epithelial loss of *Bmal1* affects fucosylated and sialylated mucus production. (**a**)Representative images of *Epi^Ctrl^* and *Epi^ΔBmal1^* distal colon stained for UEA-1 (violet) and MALII (yellow) to label fucosylated and sialylated mucins, respectively. Shown are samples collected at ZT0 and ZT12, arrowheads indicate mucus thickness. (Scale bar = 20μm). (**b**) UEA-1+ mucus thickness in the distal colon is significantly higher at ZT12 in *Epi^Ctrl^* but lower at both times in *Epi^ΔBmal1^* (**p≤0.01). (**c**) MALII+ mucus thickness in the distal colon is significantly higher at ZT12 in *Epi^Ctrl^* (*p≤ 0.05), with a significant increase of thickness in *Epi^ΔBmal1^* at ZT0 (*p<0.05). Note, that *Epi^ΔBmal1^* does not show a time-of-day dependent change in MALII. (**d**) UEA-1+ goblet cell area in distal colon crypts s significantly higher at ZT12 in *Epi^Ctrl^* (**p≤0.05). While not significantly different from controls, *Epi^ΔBmal1^* do not show this time-of-day change in UEA-1+ mucus thickness. (**e**) MALII+ goblet cell area in distal colon crypts is significantly increased in *Epi^ΔBmal1^* at ZT0 (*p≤0.05).

### Staining and histology

Fixed colon tissues were washed 3 times in PBS to remove any PFA before being submerged in 30% (w/v) sucrose in PBS for 24 hours at 4°C. Tissue was mounted in Tissue-Tek O.C.T compound (Supp. akura, Flemingweg, Netherlands) and left at 4°C for 1 hour before being frozen at -80°C. The frozen tissue was sectioned at 10μm using a Leica CM 1950 Cryostat (Leica Biosystems). Tissue morphology was assessed using Hematoxylin & Eosin (Electron Microscopy Sciences, Hatfield, PA). DAB staining was performed using HRP/DAB (ABC) Detection IHC Kit as per manufacturer’s protocol (ab64261, Abcam). When necessary, antigen retrieval was done using a 1x sodium citrate buffer. Antibodies and staining protocol used are: anti-Ki67 (ab16667, Abcam), anti-pHH3 (06-570, Upstate/ Millipore), anti-Villin (ab130751, Abcam). Hematoxylin & Eosin (H&E), Alcian blue, and Periodic Acid Schiff (PAS) staining was performed as previously described (21). Alcian blue, PAS, and Lectin staining were performed on Carnoy’s-fixed paraffin-embedded colon sections. For Lectin staining, slides were deparaffinized in xylene, rehydrated through a graded ethanol series, and blocked for endogenous biotin/streptavidin binding using a Streptavidin/Biotin blocking kit (Vector Laboratories). Sections were incubated with biotinylated MAL-II (B-1265-1, Vector Laboratories; 3µg/mL in 1%BSA in PBS) overnight at 4°C, washed in PBS, then incubated with UEA-1 rhodamine (RL-1062-2, Vector Laboratories; 2µg/mL) and streptavidin-FITC (405201, BioLegend; 5µg/mL) in 1% BSA in PBS for 1 hr at room temperature in the dark. RNA fluorescence in situ hybridization (RNA-FISH) was performed by first post-fixing slides in 4% paraformaldehyde (PFA). RNA-FISH was performed according to the HCR™ RNA-FISH protocols developed by Molecular Instruments for formalin-fixed paraffin-embedded (FFPE) tissue sections (Revision 4). In the FFPE protocol, a 5-minute incubation with 1% SDS at room temperature was performed in place of Tris-EDTA and proteinase K was omitted from both protocols. *16S rRNA* was detected using a general “Bacillus subtillis-*EUB-338”* HCR probe.

### Microscopy and image analysis

Samples were imaged using a Zeiss Axio Scan.Z1 slide scanner or Zeiss LSM900 Confocal (Zeiss, Toronto, Canada). Crypt length and area measurements were collected using the Zen Blue Software (Zeiss) and analyzed as *per* our previous studies(21). Images were processed using Adobe Photoshop (San Jose, CA). Antibody-positive cells were quantified as the total number of labeled cells/nuclei, crypt, or area. Ten crypts were sampled per animal for each colon region (proximal and distal) for quantification of Ki67 and pHH3. Mucus thickness was measured by sampling 15 regions per transverse cross section and averaging at least 3 cross sections of proximal and distal colon per mouse, data presented are averages of 5-6 mice per treatment. Goblet area was quantified by sampling five regions of well-oriented crypts (2-3 crypts per region) from at least three distinct cross-sectional regions per tissue section. Positive staining was outlined and quantified as area fraction per crypt using Zen Blue software (Zeiss). All measurements were averaged per mouse for statistical analysis.The distance between the epithelial barrier and the *16S rRNA* bacterial layer was measured in 40 proximal and distal regions and averaged per mouse.

### Quantitative PCR

*Bmal1* gene expression was normalize to the control gene Gapdh, and tested using iTaq Universal SYBR Green Supermix (Bio-Rad) on a Viia7 real-time polymerase chain reaction plate reader (Thermo Fisher Scientific). The primers used were *Bmal1* (forward: TGACCCTCATGGAAGGTTAGAA, reverse: GGACATTGCATTGCATGTTGG); Gapdh (forward: AGGTCGGTGTGAACGGATTTG, reverse: TGTAGACCATGTAGTTGAGGTCA).

### Fecal lipocalin analysis

Mice were individually housed in cages with foam pad bedding to facilitate daily stool collection and ensure the removal of old stool between collections. Stool samples as indicated in figures were analyzed for fecal LCN2 content using ELISA.

### Statistics and bioinformatics

Statistical analysis was performed using Prism 10 (Graphpad). Tests including Unpaired *t*-test, One-way ANOVA, or Two-way ANOVA were used with Tukey’s multiple comparisons or Šídák’s multiple comparisons post-test, respectively; see Supplementary Table 1 for all statistical details. Analysis of rhythmic genes(34) identified was done using Excel (Microsoft) and Cytoscape(35) software. Protein-protein interaction networks were established with medium confidence settings (0.400-0.700), and GO enrichment was carried out on the resulting networks.

## Results

### Epithelial loss of *Bmal1* reduces mucus barrier production

*Bmal1* is a non-redundant clock transcription factor, whose loss disrupts molecular and physiological circadian rhythms (26). We crossed *Villin-cre* mice (36), that have epithelial-specific expression in the colon (Supplementary Fig 1a), with *Bmal1^flox/flox^*mice (27) to conditionally delete *Bmal1* (Supplementary Fig 1b). We tested control *Vil^+/+^; Bmal1^flox/flox^* (*Epi^Ctrl^*) and experimental *Vil^Cre/+^; Bmal1^flox/flox1.^* (*Epi^ΔBmal1^*) for the effects of *Bmal1* loss of function in the colon epithelium. First, the daily locomotor activity of mice was monitored. Daily rhythms persist in both *Epi^Ctrl^* and *Epi^ΔBmal1^*, with increased activity taking place during the dark phase (Fig 1a-b). Overall, no significant differences in day *vs*. night activity between the two genotypes was observed (Fig 1c), these display the same rhythmic amplitude and mesor (Fig 1d). These results, which are consistent with previous work (31, 32), demonstrate that the loss of *Bmal1* in the epithelium does not affect behaviour rhythms.

Previous studies that have tested epithelial *Bmal1* loss of function in the context of colitis did not report if differences in undamaged colon tissues are present in undamaged pre-colitis conditions (29, 31–33). Careful examination of tissue sections in the proximal and distal regions of the colon revealed a morphology generally indistinguishable from controls (Fig 1e-h), however, unlike the *Bmal1^-/-^* mutant the *Epi^ΔBmal1^* colon displays an increased density of crypts in the distal colon (Fig 1h). This increased cellular density suggests a slight disruption in tissue architecture that occurs when *Bmal1* is lost in epithelial cells.

Mucus is a barrier in the gastrointestinal tract, where layers of mucus secreted in the proximal *vs.* distal colon goblet cells contribute to microbiota resistance (12). To determine if epithelial *Bmal1* is associated with barrier function, we performed an analysis of mucus levels. We used Carnoy’s fixation to preserve mucus integrity, and stained undamaged *Epi^Ctrl^* and *Epi^ΔBmal1^* colons with Alcian blue staining (Fig 2a-c), a marker of acidic mucus (37), and Periodic Acid Schiff (PAS), a marker of neutral and acidic mucus (37) (Fig 2d-e). In the *Epi^Ctrl^* distal colon, Alcian blue shows a time-dependent change in thickness, with higher levels at ZT12 than at ZT0 (Fig 2a-b). This thickness is disrupted in the *Epi^ΔBmal1^* distal colon, where mucus thickness was significantly lower at both timepoints relative to controls. To determine if more mucus is produced at ZT12, the Alcian blue area in goblet cells in colon crypts was quantified. This revealed a significant increase at ZT12 compared with ZT0 in *Epi^Ctrl^*, along with a significant decrease in *Epi^ΔBmal1^* at both timepoints (Fig 2c). PAS staining confirmed the changes in mucus thickness observed in Alcian blue, with similar increases at ZT12 and a reduction in *Epi^ΔBmal1^* (Fig 2d-e). Together these data suggest that more mucus is produced when mice become active (ZT12) than when they are inactive (ZT0), and that this production is in part regulated by *Bmal1* in colonic epithelial cells. Of note, *Epi^ΔBmal1^* show a slight but significant increase in mucus at ZT12 like the control (Fig 2b-c, 2e), indicating that epithelial clock function is not the only regulator of a time-dependent change in mucus levels.

We next examined whether the decreases in mucus thickness in *Epi^ΔBmal1^* mice impact barrier function. Staining for bacterial 16S RNA at ZT0, following nocturnal feeding when mice are active, showed that distal colon bacteria in *Epi^ΔBmal1^* are indeed physically closer to epithelial cells (Fig 2f-g). We tested feces of *Epi^Ctrl^* and *Epi^ΔBmal1^* mice for the levels of Lipocalin, a marker of intestinal inflammation, to see if this increased bacterial proximity leads to an increase in inflammatory responses; *Epi^ΔBmal1^*mice have no significant increase in Lipocalin though we do note a trend to increased levels (Fig 2h, p=0.06).

Given the decrease in mucus production, we further probed for *Ulex europaeus* agglutinin I (UEA-1), a fucose-binding lectin that stains mucus enriched in upper crypt and surface Goblet cells (38), and *Maackia amurensis* lectin II (MALII), a sialylated and sulfated glycan binding lectin that marks mucus in the distal mouse colon (39). UEA-1+ mucus marks both barrier and niche mucus layers, while MALII mucus is thought to be a germ-free barrier that prevents microbiota contact with epithelia; both typs of mucus are present in *Epi^Ctrl^* and *Epi^ΔBmal1^*mice. In the *Epi^Ctrl^* distal colon, UEA-1+ mucus shows a time-dependent change in thickness, higher at ZT12 than at ZT0, like the Alcian blue and PAS staining, and this pattern is lower in Epi *^ΔBmal1^*at both timepoints (Fig 3a-b). MALII+ mucus shows a time-dependent change in mucosal thickness in the *Epi^Ctrl^*distal colon, with higher thickness at ZT12 than ZT0. In contrast, Epi *^ΔBmal1^*distal colons have significantly thicker MALII+ mucus relative to controls, with a loss of time-dependent variation (Fig 3a, c). To determine if more fucosylated mucus was produced at ZT12, the UEA-1+ area in the goblet cells of distal colon crypts was quantified. This revealed a significant increase at ZT12 in *Epi^Ctrl^*, with no time-dependent difference in Epi *^ΔBmal1^* colons, suggesting that temporal control of fucosylated mucus requires epithelial *Bmal1* (Fig 3d). To determine the same for sialylated mucus, the MALII+ area in the goblet cells of distal colon crypts was quantified. A significant increase in the MALII+ area was observed in the Epi *^ΔBmal1^*colon at ZT0 (Fig 3e). *Epi^Ctrl^*colons show no time-dependent variation in MAL-II+ goblet area, although a trend was observed (p=0.0503).

We further tested UEA-1+ mucus in proximal goblet cells to determine whether these contributes to the distal mucus barrier thickness. This revealed a significant decrease in *Epi ^ΔBmal1^* (Supplementary Fig 2). No time-dependent variation was observed in *Epi^Ctrl^* proximal colons, suggesting that temporal control of mucus production is distal colon specific. Of note, a sex-specific difference was observed, with Epi *^ΔBmal1^* males having a slight reduction in proximally derived UEA-1+ mucus (Supplementary Fig 2c). Altogether, these results suggest that mucus thickness, and its fucosylation and sialylation, are time-dependent with an increase towards the active phase (ZT12). The loss of mucus in Epi*^ΔBmal1^*, suggests that *Bmal1* is, at least in part, required for mucus production.

### Epithelial *Bmal1* loss produces a male-specific increase in colitis

Our previous studies of the *Bmal1* null mutant showed greater inflammation when mice are treated with Dextran Sulfate Sodium (DSS), an injury-based model of colitis (21). Several publications have reported test of the *Villin-cre* + *Bmal1* epithelium conditional knockout in the context of DSS-colitis, however, these have been done only using male mice (30, 32, 33). We thus asked whether a sex-specific effect of *Epi^ΔBmal1^* mice is a factor. To test this, a cohort of male *vs.* female *Epi^Ctrl^* and *Epi^ΔBmal1^* mice of ∼4 months of age were housed in individual cages, treated with 2.5% DSS for seven days, and then monitored for recovery, weight loss, and disease activity (Fig 4a). Male *Epi^ΔBmal1^* mice showed significantly poorer recovery from colitis compared to female mice (Fig 4b), as well as an increase in weight loss (Fig 4c), in line with a previous report that shows this strain to be more susceptible to colitis (30). However, we did not find an increase in disease symptoms relative to controls which is consistent with a different report (32) (Fig 4d). We did not observe that *Epi^ΔBmal1^* male show improved resistance to colitis, in contrast to another study (33). These effects were sex-specific: *Epi^ΔBmal1^* female mice are indistinguishable from controls (Fig 4b-d). To further test inflammation during colitis and recovery, we again tested feces of *Epi^Ctrl^* and *Epi^ΔBmal1^* mice for Lipocalin. The application of DSS greatly increases the levels of this inflammatory marker as would be expected, which then decrease during regeneration post-DSS withdrawal (Supplementary Fig 2a-b). No significant differences between *Epi^Ctrl^* and *Epi^ΔBmal1^* Lipocalin are evident over the course of the experiment. Of note, the mice that were humanely euthanized due to poor DSS recovery (Fig 5b) would not have had their feces examined from that timepoint onward, so these data likely underestimate Lipocalin levels in *Epi^ΔBmal1^* male mice. Together, these data indicate there is a male-specific *Bmal1* epithelial loss of function phenotype in the context of DSS-induced colitis.

**Figure 4:**
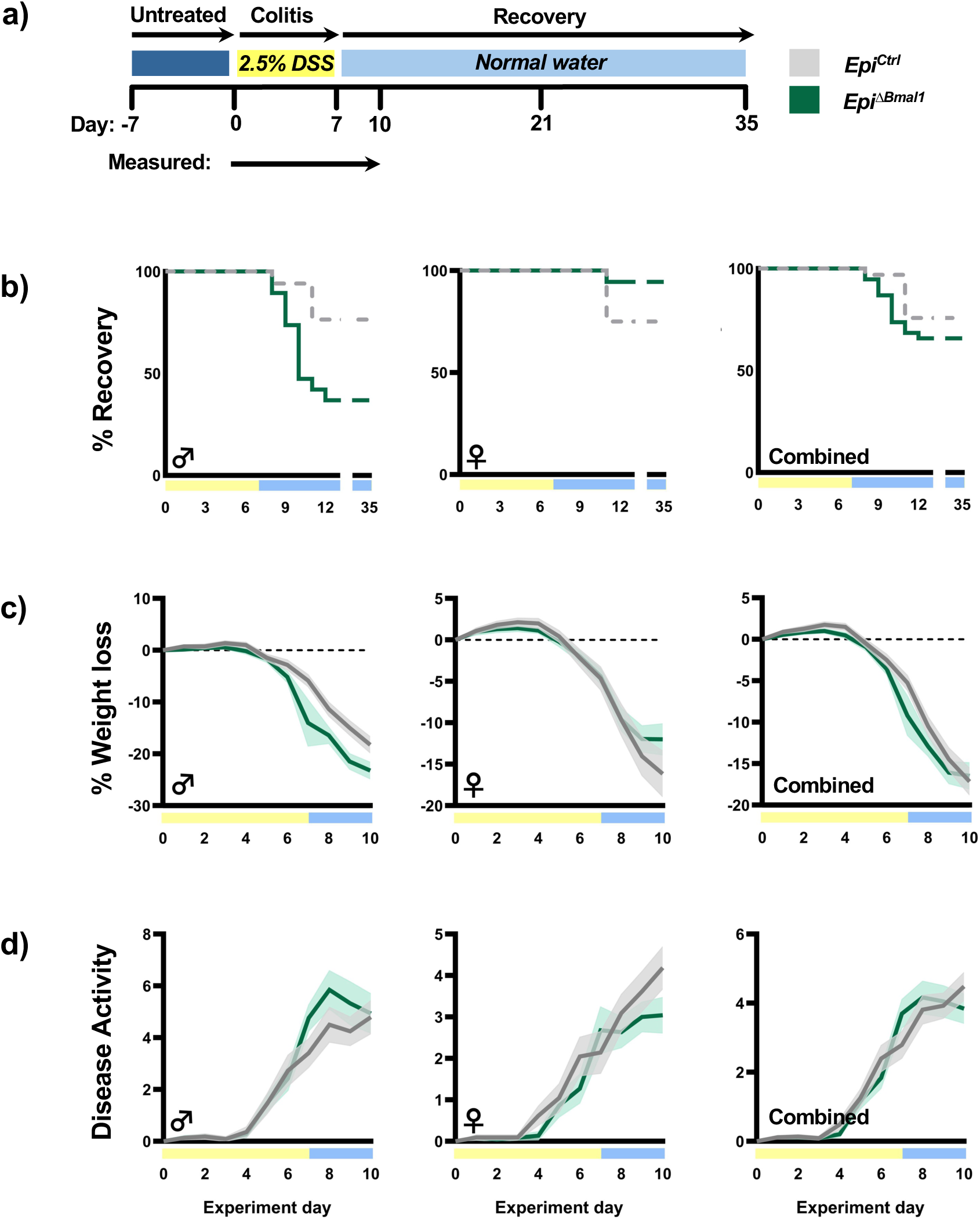
Epithelial *Bmal1* loss produces a male-specific increase in colitis. (**a**) Experimental schematic demonstrating DSS administration and recovery of *Epi^Ctrl^* and *Epi^ΔBmal1^*. (**b**) Recovery of mice following DSS reveals that male (♂) *Epi^ΔBmal1^*mice have a decreased ability to recover (Mantel-Cox test, *p ≤ 0.05); females (♀) show no significant differences; recovery from males and females combined shows no significant differences. (**c**) Percent of total weight loss is significantly decreased in ♂ *Epi^ΔBmal1^* mice (Two-way ANOVA, ***p ≤ 0.001, day 7 ***p ≤ 0.001, day 9 *p ≤ 0.05); (f) ♀ do not have significant differences in weight loss; data from males and females combined shows no significant differences. (**d**) Disease Activity does not differ between *Epi^Ctrl^*and *Epi^ΔBmal1^* ♂, ♀, or combined (no significant differences). Full statistical details are provided in Supplementary Table 1.

**Figure 5:**
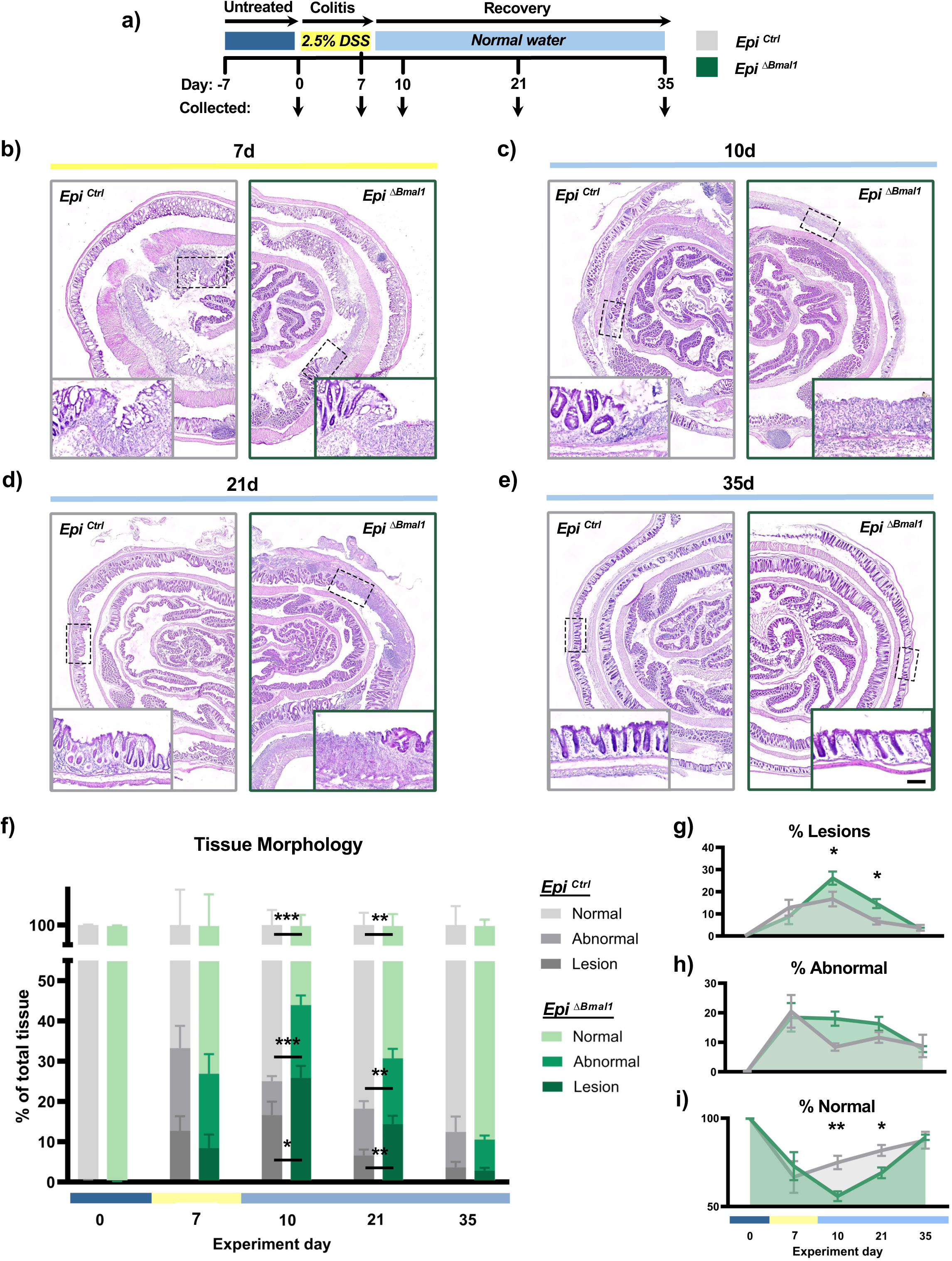
Epithelial *Bmal1* loss increases colon damage. (**a**) Experimental schematic demonstrating DSS administration and recovery of *Epi^Ctrl^* and *Epi^ΔBmal1^*. As indicated, samples were collected at day 0 (undamaged), day 7 (last day of DSS), day 10/21/35 (post-DSS withdrawal). (**b-e**) Representative images of H&E-stained colon prepared as Swiss rolls and sectioned transversally from *Epi^Ctrl^* and *Epi^ΔBmal1^* mice. Time of sample collection is indicated; magnified inset is indicated by a dotted outline highlighting the selected region shown (scale bar= 50μm). (**f**) Stacked graph shows the percentage of total colon tissue made up of normal regions (regular crypt architecture), lesions (absence of architecture), abnormal regions (regenerating hyperplastic crypts). These correspond to the images shown in (b-e). *Epi^ΔBmal1^* mice at day 10 have an increase in lesions (Unpaired t-test, *p ≤ 0.05), abnormal tissue (***p ≤ 0.001), and a decrease in overall normal tissue (***p ≤ 0.001). *Epi^ΔBmal1^* mice at experimental day 21 have a similar increase in lesions (Unpaired t-test, **p ≤ 0.01), abnormal tissue (**p ≤ 0.01), and a decrease in overall normal tissue (**p ≤ 0.01). (**g**) *Epi^ΔBmal1^* show increased lesions at days 10 and 21 (Two-way ANOVA, *p ≤ 0.05). (**h**) Abnormal regions show no significant differences in *Epi^Ctrl^*and *Epi^ΔBmal1^* mice. (**i**) A decrease in *Epi^ΔBmal1^*normal tissue is present at days 10 and 21 (Two-way ANOVA, *p ≤ 0.05). Full statistical details are provided in Supplementary Table 1.

### Epithelial *Bmal1* loss increases colon damage

In order to better understand the effects of *Epi^ΔBmal1^* loss of function, we investigated the recovery phase post-colitis when the tissue regenerates to return to its homeostatic set point. We predicted DSS-colitis injury would perturb the *Epi^ΔBmal1^*colon, revealing disrupted regeneration. Mouse proximal and distal colon tissues were analyzed pre- and post-DSS administration (Fig 5a). Tissue status was categorized by morphology: 1) regions with regular crypt architecture was categorized as “normal”; 2) regions with severe damage lacking any crypt architecture was categorized as “lesion”; 3) regions consisting of hyperplastic and/or bifurcated crypts undergoing active regeneration was categorized as “abnormal”. Initially, both the *Epi^Ctrl^* and *Epi^ΔBmal1^* mice show completely normal histology (Fig. 1e). During colitis (DSS treatment at day 7), both genotypes exhibit a similar increase in lesions and abnormal crypts (Fig. 5b). The majority of lesions are located in the middle to distal areas of the colon while the proximal regions remained largely unaffected. Abnormal areas representing regeneration were noted typically at the periphery of lesions, indicating regions that undergo regeneration subsequent to damage. Upon DSS removal, a combination of normal, lesions, and abnormal tissue remains present in both genotypes (days 10 and 21 of the experiment) (Fig. 5c-d). Finally, at later stages (experimental day 35), both *Epi^Ctrl^* and *Epi^ΔBmal1^* colons return to normal tissue architecture with few remaining lesions and abnormal regions present (Fig 5e). These morphological changes and timing of colon tissue regeneration following DSS-injury is consistent with previous studies (15, 17, 18). We also measured colon length as a marker of inflammation in these samples. While DSS significantly decreased the length of colons of both *Epi^Ctrl^* and *Epi^ΔBmal1^* at days 7, 10, and 21, we noted no differences between these genotypes (Supplementary Fig 3a-c).

We calculated each tissue morphology category as a percentage of the entire length of the colon in tissue Swiss rolls (Fig 5b-e). This analysis revealed that *Epi^ΔBmal1^* exhibit significantly more severe lesions and abnormal areas at day 10 and day 21, indicating that damage and subsequent regeneration is higher in the absence of epithelial *Bmal1* (Fig 5f). This is evident when lesion measurements are replotted individually over the course of injury/recovery, *Epi^ΔBmal1^* mice have increased damage after DSS-treatment when compared to *Epi^Ctrl^* mice (Fig 5g). This is accompanied by higher levels of abnormal (regenerating) tissue at these times, concomitant with lower percentages of normal regions (Fig 5h-i). Of note, no sex-specific differences in these changes or timing were observed. To confirm this increase in regeneration at 21 days, we compared the hyperplastic, proliferating (Ki67+ and pHH3+) crypts in *Epi^Ctrl^* and *Epi^ΔBmal1^*, which revealed regenerating tissue is indeed increased in *Epi^ΔBmal1^* (Supplementary Fig 4a-d). Together, these data indicate that *Epi^ΔBmal1^* colons show increased damage and delayed recovery compared to *Epi^Ctrl^*.

## Discussion

Our study adds to a growing area of research on the physiological function of the circadian clock, regulated by *Bmal1*, in colon epithelial cells. We find that the mucus barrier layer in the colon shows time-dependent changes in thickness, higher when mice become behaviourally active and start feeding, than when they are inactive (Fig 2-3). This change is linked to *Bmal1* function in colon goblet cells, *Epi^ΔBmal1^*mice have lower mucus production (Fig 2-3). We also find that fucosylation and sialylation of mucus mirrors these changes, with the *Epi^ΔBmal1^* epithelial loss of function showing disruptions in the timing of these. While the absence of epithelial *Bmal1* is not detrimental in mouse laboratory conditions, the colon exhibits a phenotype when *Epi^ΔBmal1^* mice are challenged by injury. Epithelial *Bmal1* loss of function leads to a male-specific outcome, with males showing reduced recovery during DSS-colitis (Fig 4). Long-term outcomes of the loss of epithelial *Bmal1* are also apparent: it is required during regeneration to promote the efficient restoration of normal tissue architecture post-injury in both males and females (Fig 5). Our results highlighting the importance of *Bmal1* in the colon epithelium are broadly consistent with previous studies (29, 30, 32), and our work corroborates the male-specific susceptibility of one of these studies (29). We extend these previous reports to show that *Bmal1* is also important in regeneration and homeostatic recovery following colitis. Taken together, it becomes clear that there are several possible mechanisms of how the circadian gene, *Bmal1*, aids in the maintenance of the colon barrier during homeostasis.

An important function of *Bmal1* in the colon epithelium is to regulate either the rhythmicity or composition of the microbiome (29–32). The loss of epithelial *Bmal1* is proposed to have pathological consequences including increases in pro-inflammatory bacteria (29, 30), decreases in barrier function (29), and increases in inflammatory responses (29, 30, 32). Our findings that mucus function is *Bmal1*-dependent, supports the possibility that a lowered mucus barrier may be upstream of cellular responses to microbiota (13, 14), subsequent environmental challenge including with DSS would then activate the immune system to drive increased inflammation. This would explain why *Bmal1* conditional epithelium knockouts generally display increased colitis phenotypes. The DSS-injury model is microbiome-dependent (40–42), and it is likely that differences in microbiota composition in the colon are present in the different labs that have tested the *Bmal1*-epithelial loss of function. A thinner mucus barrier coupled with variability in microbiota populations would parsimoniously explain why some labs have noted variability in colitis outcomes in the epithelial *Bmal1* DSS assays.

Mucosal modifications such as fucosylation and sialylation are central to barrier structure and integrity. Fucosylation, a process whereby fucose residues are attached to glycans by Fucosyltransferase 2 (Fut2), promotes bacterial tolerance (43). Mutations in *FUT2* are associated with increased susceptibility to IBD in patients (44), and in mouse models, an epithelial-specific knockout of *Fut2* results in exacerbation of DSS-induced colitis (45). Sialylation is a process in which sialic acid residues stabilize mucins, and this deficiency also results in increased colitis in IBD patients (46) and mouse models (47, 48). Our findings indicate that mucin fucosylation and sialylation are temporally regulated, with *Bmal1* playing a role in regulating their daily timing.

The conditional deletion of *Bmal1* in the colon epithelium differs from studies examining circadian clock mutants in colitis. When *Bmal1* is genetically disrupted only in the epithelium, its function, including its role in driving circadian transcriptional rhythms, is abolished in a subset of colon cells. Yet the surrounding cells in the colon have rhythms, as do those in the rest of the body. On the other hand, a clock gene mutant abolishes function globally, not only in the epithelial cells. Hence, the *Bmal1* mutant or *Per1/2* double mutant that have no circadian rhythmicity, and thus in some manner resemble environmentally disrupted rhythms, a conditional loss of function is an artificial experimental context designed to investigate cell-specific functions that do not otherwise occur. Nonetheless, studies testing global clock gene loss of function have noted increased colitis (21, 23, 25) and share two important features with a *Bmal1* epithelial loss of function studies (29, 30, 32). First, these studies all note that clock gene loss has a mild phenotype in the colon, that is exacerbated during DSS-colitis due to excessive inflammatory signaling through IL1β / IL6, and/or TNF / NFκB pathways (21, 23, 25). This is consistent with both our findings and studies investigating *Bmal1* in the epithelium during DSS-colitis (29, 30, 32). Of note, a *Rorα* epithelial loss of function has also been carried out that shows increased colitis due to upregulation of inflammatory NFκB signaling (22). Second, it has been noted that the clock disrupted mice have defects in epithelial proliferation and cell cycle control (21, 23). This is consistent with our findings that the epithelial *Bmal1* loss of function has delayed regeneration. These pro-inflammatory and regeneration-deficient phenotypes are consistent with our present work, and that of other groups testing the *Bmal1* epithelial loss of function (29, 30, 32).

Further supporting the role of the clock in inflammation, a 2018 study of the *Nr1d1* gene revealed it represses the Nlrp3 inflammasome and that the *Nr1d1* mutant has increased colitis (25). The Nlrp3 inflammasome is a sensor of bacteria that initiates immune system responses, which has been implicated in regulating inflammatory circadian rhythms in macrophages (49–51). Disruptions to this pathway in mouse models is associated with inflammatory and autoinflammatory phenotypes including colitis. In the epithelium, the inflammasome gene, *Nlrp6*, is highly expressed in goblet cells where it activates mucus production in response to the detection of potential pathogens (52–55). These studies raise the question of whether the thickness of the mucus barrier precedes the detection of bacteria, or whether an initial defect in bacterial detection causes a reduction in mucus thickness. Future work in this area will resolve which of these mechanisms is controlled by the circadian clock in goblet cells.

We note that our findings, and those of others (29, 30), do not support a role for *Bmal1* in promoting colitis, in contrast to Hua *et al.* (2025) (33), where a *Bmal1* epithelial loss of function led to a reduction in DSS-colitis symptoms. In this study, it was found that *Bmal1* transactivates the expression of pro-apoptotic genes (*p53*, *Bax*, *Bak1*) which increase cell death in a time-dependent manner in control mice (33). The five studies have certain differences in methodology which should be noted in light of this discrepancy. The studies demonstrating increased colitis in epithelial *Bmal1* loss, used either a *Villin-cre* or *TS4-cre* which are expressed during pre- and post-natal gastrointestinal epithelium that remains high in the adult. Hua *et al.* used a *Villin-creERT* driver that was induced only in the adult by tamoxifen injection, this may mean that early loss of *Bmal1* contributes to the opposite colitis phenotype due to an unappreciated developmental effect. However, we do not think a developmental effect is likely to be a primary factor. The gastrointestinal tract is essentially normal in the *Epi^ΔBmal1^*mouse histologically (Fig1, Supplementary Fig 4), and no inflammation is observed until colitis is induced by DSS. No pre- colitis phenotypes have been reported by other *Bmal1* conditional loss of epithelium function studies (29–32), and notably other clock mutants also show normal histology and do not have inflammation until the induction of colitis by DSS (21–25). The levels of DSS used in our study and others are similar (2.0-2.5%) and the conditional *flox* allele to disrupt *Bmal1* is the same. We thus favour the interpretation that a microbiome-dependent or mucus-dependent effect contributes to the discrepancies between studies, which (despite their discrepancies) broadly agree on the importance of the clock in regulating a time-dependent colitis phenotype.

The circadian clock has emerged as a regulator of both tissue physiology, immunity, and microbiota rhythms. It is perhaps not surprising that in the context of colitis, a disease underpinned by these factors, the clock plays a critical role. Our work and that of others highlights how the loss of circadian clock rhythms is detrimental, and further supports future work targeting the circadian system to prevent and/or treat disease.

## Acknowledgements

We are grateful to technical staff at the Universities of Windsor and McMaster University, and Kirk Bergstrom (University of British Columbia) for their help and advice. To the authors whose research we could not cite due to space limitations, we offer our apologies and thanks.

## Author Contributions

Conceptualization and experimental design was done by ZT, CE, PK; data collection was done by ZT, CE, RK, MF, MH, and HW; data analysis was done by ZT, CE, PK; writing was done by ZT, CE, WK, and PK; supervision and project administration was done by WK, and PK. All authors have read and agreed to the final version of the manuscript.

## Supplementary Material

**Supplementary Figure 1:**
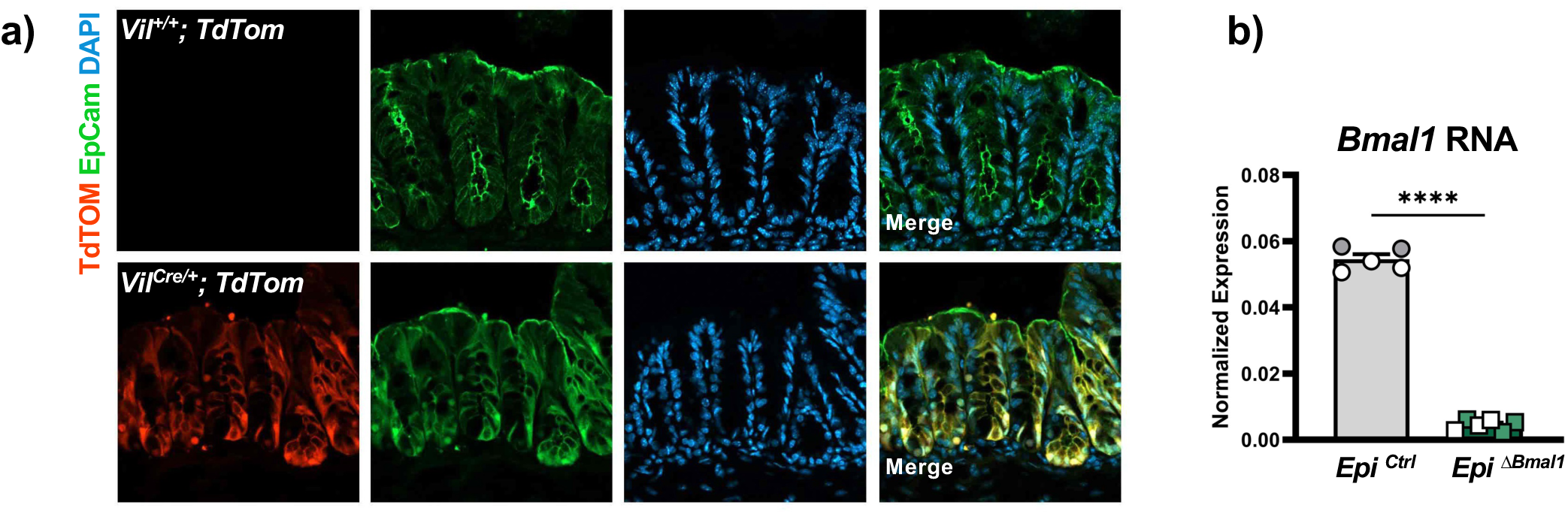
Validation of the epithelial *Bmal1* conditional knockout. (**a**) Representative image showing tdTom (red) expression in *Villin*-expressing epithelial cells only when Cre-recombinase is present in *Vil^Cre/+^;TdTom* mice (Epcam+, green) (Scale bar= 50μm). (**b**) RT-qPCR of colon tissue shows a significant decrease in *Bmal1* expression in *Epi^ΔBmal1^* mice compared to *Epi^Ctrl^* (Unpaired t-test, ****p ≤ 0.00001). In the graphs, male mice are indicated in white symbols and female mice as coloured (grey or green) symbols. Full statistical details are provided in Supplementary Table 1.

**Supplementary Figure 2:**
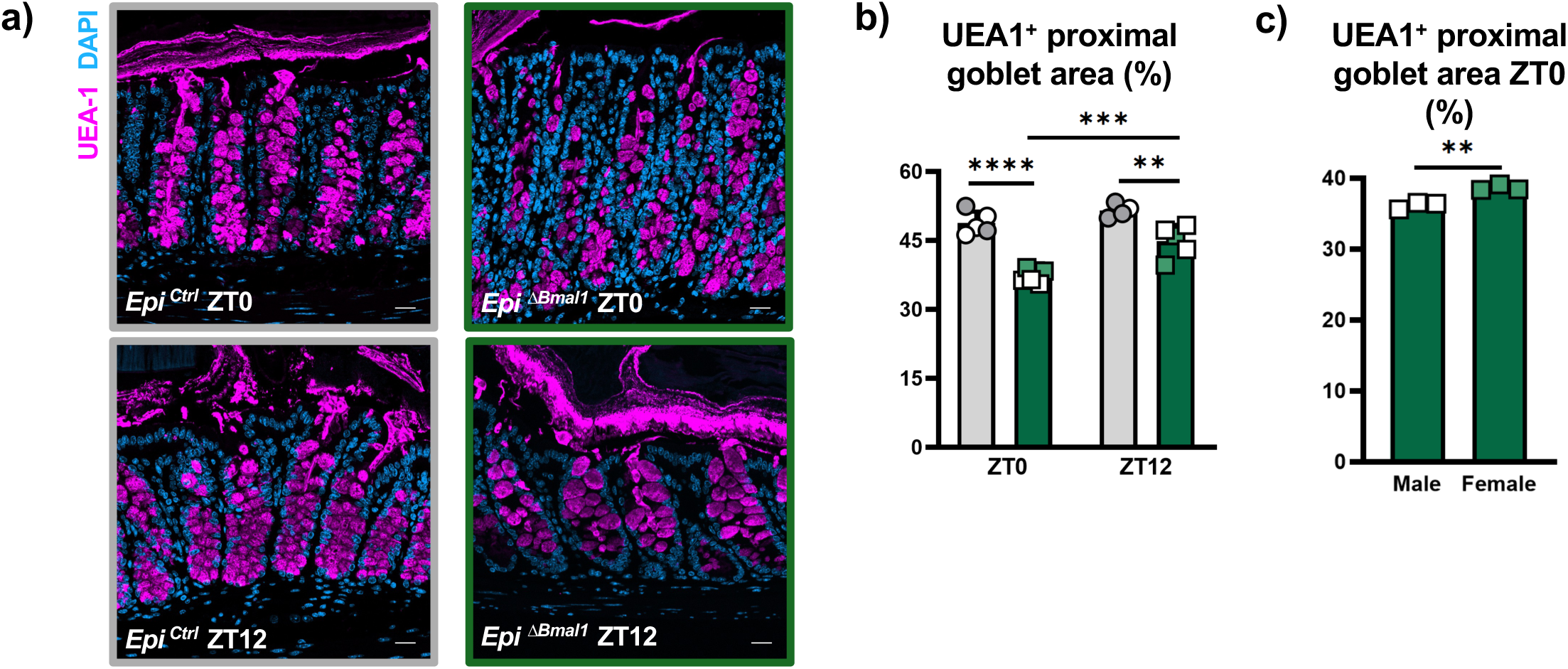
Fucosylated mucus is reduced in *Epi^ΔBmal1^*mice in the proximal colon. (**a**) Representative images of *Epi^Ctrl^* and *Epi^ΔBmal1^* proximal colon stained for UEA-1 (violet), at ZT0 and ZT12. (Scale bar = 20μm). (**b**) UEA-1+ goblet cell area in proximal colon crypts is significantly decreased in *Epi^ΔBmal1^* at ZT0 (****p≤0.0001) and ZT12 (**p≤0.01). *Epi^ΔBmal1^* has significantly increased area at ZT12 compared to ZT0 (***p≤0.001). (**c**) Goblet cell area is decreased in *Epi^ΔBmal1^* males at ZT0 (**p≤0.01). No other sex-linked effects were noted.

**Supplementary Figure 3:**
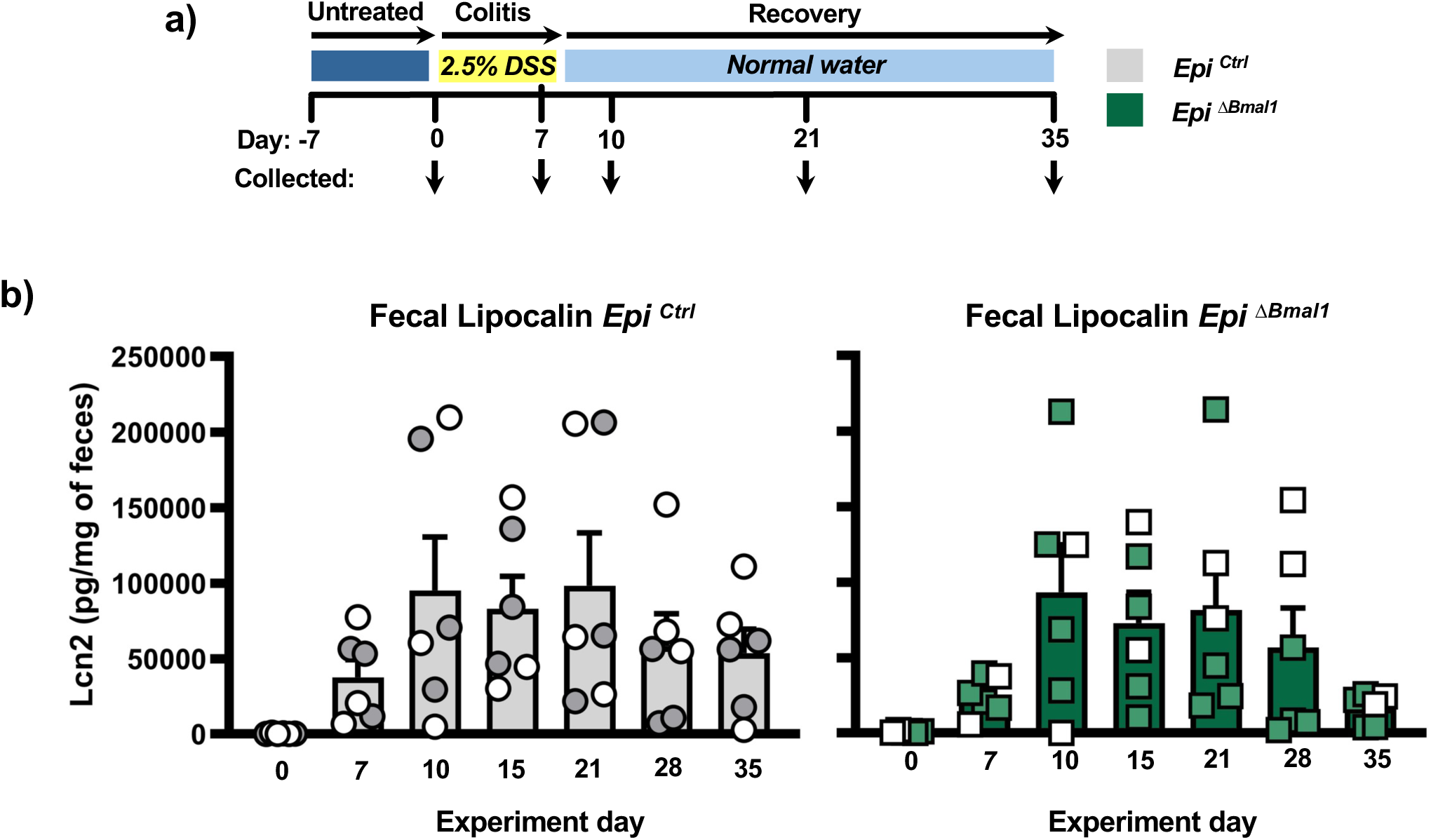
Fecal lipocalin is not affected by epithelial *Bmal1* loss during colitis and recovery. (**a**) Experimental schematic demonstrating DSS administration and recovery of *Epi^Ctrl^* and *Epi^ΔBmal1^*. (**b**) Graphs showing lipocalin (Lcn2) levels in feces in pg detected *per* mg. Initial levels are low and increase following DSS application (withdrawn on day 7) to reach a maximum at day 10 – day 21. Thereafter levels are reduced. There are no significant differences between genotypes observed. In the graphs, male mice are indicated in white symbols and female mice as coloured (grey or green) symbols. Full statistical details are provided in Supplementary Table 1.

**Supplementary Figure 4:**
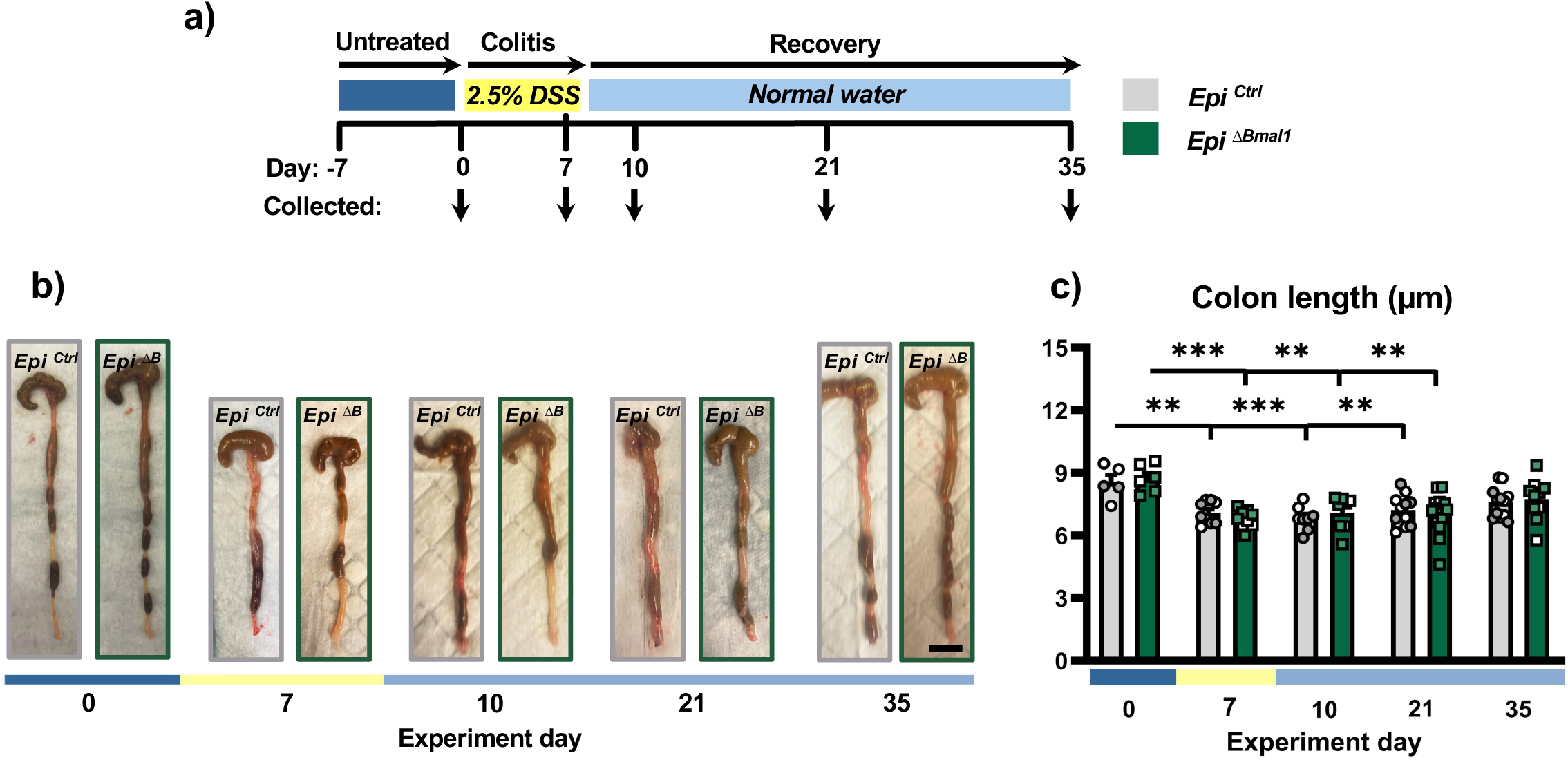
*Epi^ΔBmal1^* colon length is equivalent to controls. (**a**) Experimental schematic demonstrating DSS administration and recovery of *Epi^Ctrl^* and *Epi^ΔBmal1^*. As indicated, samples were collected at day 0 (undamaged), day 7 (last day of DSS), day 10/21/35 (post-DSS withdrawal). (**b**) Representative images show colon samples collected (scale bar= 1cm). (**c**) Average colon length decreases in size in during DSS-administration at day 7, day 10, and day 21 in both *Epi^Ctrl^* and *Epi^ΔBmal1^* mice, before returning to a normal length at day 35. (Two-way ANOVA, no significant differences; One-way ANOVA *Epi^Ctrl^* / *Epi^ΔBmal1^*, **p ≤ 0.01, ***p ≤ 0.001). In the graphs, male mice are indicated as white dot symbols and female mice are indicated as coloured (grey or green) dot symbols. In the graphs, male mice are indicated in white symbols and female mice as coloured (grey or green) symbols. Full statistical details are provided in Supplementary Table 1.

**Supplementary Figure 5:**
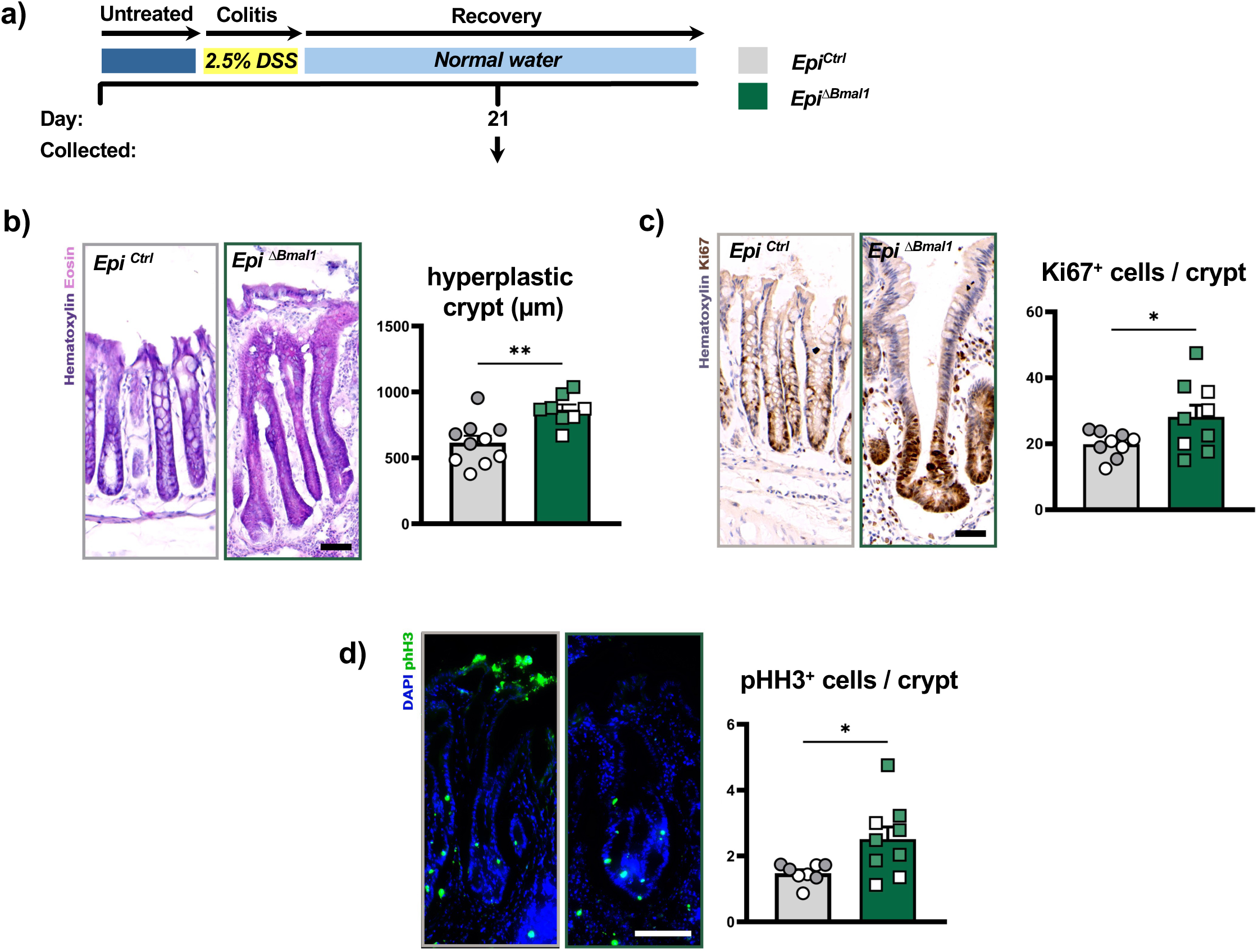
Persistent crypt proliferation in the *Epi^ΔBmal1^* colon epithelium two weeks after DSS withdrawal. (a) Experimental schematic demonstrating *Epi^Ctrl^* and *Epi^ΔBmal1^* colon samples collected on day 21. (b) Representative image of H&E-stained hyperplastic crypts from colons collected at experimental day 21 (scale bar= 50μm). The average length of hyperplastic crypts in *Epi^ΔBmal1^* colons is increased. (Unpaired t-test, **p ≤ 0.01). (**c**) Representative image of Ki67+ cells (brown) in the hyperplastic crypts of colons at day 21. The number of Ki67+ cells in hyperplastic *Epi^ΔBmal1^* crypts are higher (Unpaired t-test, *p ≤ 0.05). (**d**) Representative image of phosphorylated Histone H3 (pHH3) positive cells (green) in hyperplastic crypts at day 21 (DAPI nuclei in blue). Similar to Ki67, the number of pHH3+ cells are increased in *Epi^ΔBmal1^*(Unpaired t-test, *p ≤ 0.05). These data show that at this time, *Epi^ΔBmal1^* regeneration is delayed relative to *Epi^Ctrl^*. In the graphs, male mice are indicated in white symbols and female mice as coloured (grey or green) symbols. Full statistical details are provided in Supplementary Table 1.

